# Difference in Kernel Shape and Endocarp Anatomy Promote Dehiscence in Pistachio Endocarp

**DOI:** 10.1101/2023.08.18.553428

**Authors:** Shuxiao Zhang, Minmin Wang, Alisa Chernikova, Shaina Eagle, Kaleigh Marie Bedell, Karen Nguyen, Barbara Blanco-Ulate, Judy Jernstedt, Georgia Drakakaki

## Abstract

A fully split shell in pistachio (*Pistacia vera*) is a trait that is preferred by consumers and is a criterion in evaluating the grade of the pistachio nut. However, while the expanding kernel has been hypothesized to provide the physical force needed for shell split, the mechanisms that control shell split remains unknown. Furthermore, it is intriguing how the shell, or endocarp, splits at the suture ridge when there is no clear dehiscence zone.

The objectives of this study were to 1) identify traits associated with dehiscence in fruit in the high split rate cultivar Golden Hills when compared to the lower split rate cultivar Kerman and determine the anatomical features associated with endocarp dehiscence at the suture region, and 2) examine the effect of kernel shape on endocarp dehiscence.

We determined that despite of the fact that the pistachio endocarp is primarily composed of a single type of polylobate schlerenchyma cell, specialization of cell shape and size at the suture site results in smaller, more flattened cells. We report that there is a furrowing of the shell at the dorsal and apical suture site, where dehiscence initiates. This furrowing is not observed at the ventral suture site or in the indehiscent fruit of *Pistacia atlantica*, a species that has been used as rootstock for *P. vera*. In addition, the size of the kernel in the sagittal axis (the width) is strongly associated with higher split rate. Based on our results, a tentative model emerges where, in the absence of specialized cell types, cell shape modification can create an anatomically distinct region that is mechanically weak in the endocarp for the initiation of dehiscence, while the force from the width of the kernel is necessary for the shell split rate difference as observed in cultivars.

## INTRODUCTION

The pistachio (*Pistacia vera*) fruit is botanically classified as a drupe with three main components: kernel, shell, and hull (Ferguson et al. 2005; Polito and Pinney 1999). The edible kernel, contained within the shell, is the seed, while the shell is the heavily lignified endocarp. Exterior to the endocarp is a fleshy hull, comprised of the exocarp and mesocarp (Ferguson et al. 2005; Polito and Pinney 1999). Together, the endocarp, mesocarp, and exocarp form the pericarp of the fruit (Fig. 1), while the seed and endocarp form the commercial fruit product. Plant breeding efforts have focused on increasing the size and improving the quality of the seed, but also on the rate and extent of pericarp dehiscence at maturity (Kallsen et al. 2009; Kallsen and Parfitt 2017; Parfitt et al. 2007). Shell split, or endocarp dehiscence, is of particular interest, since this characteristic improves the ease of consumption and is therefore a desired trait for consumers and growers.

**Fig. 1.**
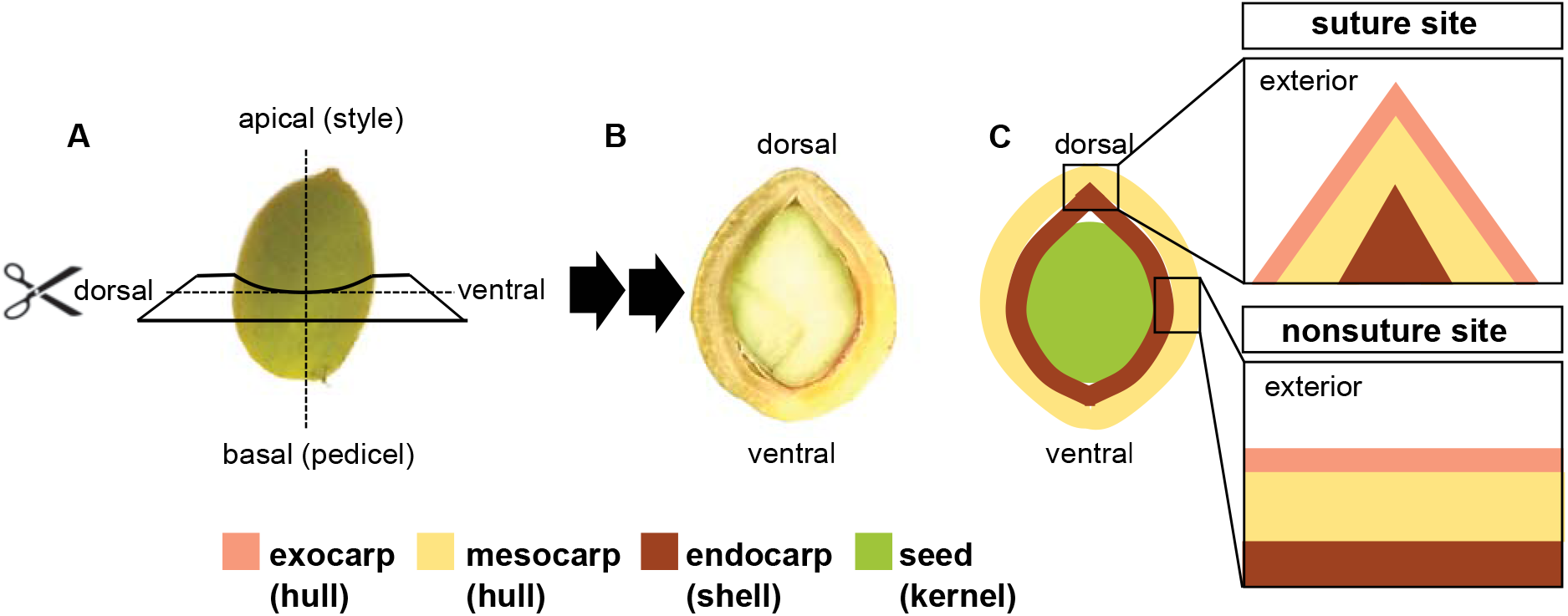
Sites sampled and organization of *Pistacia vera* pericarp. (A) Intact pistachio fruit with dashed lines indicating two planes of section, the apical-basal axis and the dorsal-ventral axis, used in anatomical analysis. The longitudinal plane runs from the apical (style) end of the fruit to the basal (pedicel) end. With the exception of the apical site, all samples were collected from the median of the fruit cut in the transverse plane as indicated by the solid lines. (B) Photograph (face up) of the pistachio fruit cut in the transverse plane indicated in panel A showing the seed (kernel) at the center surrounded by the pericarp. (C) Diagram of the pistachio fruit cut in transverse plane (face up as in B). Insets indicate regions sampled from the boxes in the larger diagram for micrographs in this study. Suture site indicates the region of initial shell split. All micrographs are presented with the exterior in the upward direction.

Fruit dehiscence has been studied in many species, such as the model plant arabidopsis (*Arabidopsis thaliana*), members of the family Fabaceae, and members of the genus *Prunus* (Dardick and Callahan 2014; Gradziel and Martínez-Gómez 2002; Han et al. 2015; Parker et al. 2020, 2021; Polito and Pinney 1999; Ogawa et al. 2009; Tani et al. 2007). In contrast to these species, pistachio fruit are morphologically different and the mechanisms for pistachio endocarp dehiscence are still largely unknown (Fabbri et al. 1998; Polito and Pinney 1999; Shuraki and Sedgley 1996a). Dehiscence of the fruit wall (the pericarp) is commonly associated with the presence of one or more dehiscence zones, which often contain specialized cells and is where cell separation occurs (Hofhuis et al. 2016; Huss et al. 2020; Ogawa et al. 2009; Parker et al. 2020, 2021). Although the pericarp of the fruit of arabidopsis and members of the Fabaceae and pistachio all naturally cracks at maturity, apparently independent of pericarp modification that occurs at germination, the pistachio is unusual in that that has an exo-mesocarp that does not split, preventing seed shed. Furthermore it does not contain a recognizable dehiscence zone, and it is currently unknown how endocarp dehiscence can occur without the presence of dedicated cell types (Ogawa et al. 2009; Polito and Pinney 1999).

Given the diversity of fruit forms in angiosperms, it can be informative to compare both evolutionarily homologous and nonhomologous fruit tissues that serve a similar function in different species. Developmentally, the endocarp of pistachio is more similar to the “pit” found in members of the genus *Prunus* than the dehiscing layers in the arabidopsis fruit. Both pistachio and *Prunus* fruit are botanically classified as fleshy drupes, and it is the endocarp that dehisces (Dardick and Callahan 2014; Ferguson et al. 2005; Tani et al. 2007). In contrast, in arabidopsis fruit (a silique), dehiscence occurs through all layers of the pericarp (Dardick and Callahan 2014; Hofhuis et al. 2016; Ogawa et al. 2009). In peach (*Prunus persica*), lignification of the endocarp, expansion of the seed, and thickness of the endocarp at the suture have all been implicated in endocarp dehiscence (Han et al. 2015; Tani et al. 2007). Although the term “suture” is commonly used to describe a ridge or indentation in the “shell” where dehiscence occurs, in almond (*Prunus dulcis*), endocarp dehiscence can occur adjacent to the suture (Gradziel and Martínez-Gómez 2002). This suggests that factors other than the presence of a suture may influence the location of endocarp dehiscence. Notably, the pistachio “shell” (endocarp) is unique in that it is composed of a single cell type, has a visible suture, and dehisces upon maturity. In contrast, other members of Anacardiaceae have previously been reported to have layered endocarp with multiple cell types (Pienaar and Van Teichman 1998). In other lineages, the lignified component of stone fruit and “nuts”, such as the endocarp of peach, almond, and pecan (*Carya illinoinensis*), seed coat of macadamia (*Macadamia* sp.), and pericarp of hazelnut (*Corylus* sp.), are composed of multiple cell types and do not dehiscence upon maturity (Gradziel and Martínez-Gómez 2002; Han et al. 2015; Huss et al. 2020; Polito and Pinney 1999; Shuraki and Sedgley 1996a; Tani et al 2007; Weis et al. 1998). It should be noted that although “nut” is generally considered to be any dry seed enclosed by a protective shell, many “nuts”, such as pistachio and almond, are not botanically considered nuts.

At the anatomical level, the endocarp of the pistachio is more similar to the shell of walnut (pericarp of *Juglans regia*), which is composed primarily of a single type of polylobate sclerenchyma cells, than to the shells of other “nuts” (Antreich et al. 2019; Polito and Pinney 1999; Shuraki and Sedgley 1996a). However, the walnut shell contains a well-defined dehiscence zone at the suture that contains flattened, unlignified cells (Huss et al. 2020). In addition, walnut does not naturally dehisce upon maturity, and prior to germination, splits only when external forces are applied, suggesting that factors other than the presence of the suture are important for the initiation of endocarp dehiscence (Huss et al 2020). Mechanical studies of the walnut shell have indicated that the three-dimensional jigsaw shape of the lobed sclerenchyma cells in the nonsuture regions of the shell and the thickness of the shell may be important for the shell’s overall strength and mechanical rigidity, while the strength of the suture zone may be important for maintaining an intact, unsplit, shell (Antreich et al. 2019; Koyuncu et al. 2004; Sideli et al. 2020; Xiao et al. 2020).

It has been hypothesized that endocarp dehiscence in pistachio, similar to that in the walnut pericarp, depends on shell thickness and the angle of the suture (Polito and Pinney 1999; Shuraki and Sedgley 1996a). However, it remains an enigma how pistachio can naturally dehisce upon maturity despite the absence of a dehiscence zone, when walnut and many *Prunus* species have a lignified “shell” with a dehiscence zone, but do not dehisce (Shuraki and Sedgley 1996a). The size of the pistachio kernel (the seed), specifically the force of the expanding kernel as it presses against the pericarp, has been implicated in endocarp dehiscence (Polito and Pinney 1999). It is currently unknown if, or to what degree shell thickness, suture angle, and kernel size contribute to the rate of shell split in pistachio, or how endocarp dehiscence occurs at the suture without a well-established dehiscence zone of specialized cells.

In this study we used two pistachio cultivars, Golden Hills and Kerman with high and low shell split rates, respectively (Kallsen et al. 2009), to analyze multiple morphological and anatomical traits hypothesized to be associated with fruit dehiscence in pistachio. We observed that the anatomy of the endocarp suture, specifically the morphology of the cells at the suture, could create a mechanically weakened “dehiscence zone” in the endocarp, and hypothesize that kernel shape may be critical for endocarp dehiscence.

## MATERIAL & METHODS

### Plant material

During bloom in 2021 in mid-Apr, 12 trees each of *P. vera* ‘Kerman’ and ‘Golden Hills’ were tagged at an orchard located at Cantua Creek, CA, USA. To avoid the effect of fruit load per branch on fruit development and phenology, only “low-load” branches, those containing one fruit cluster per branch, were used. Four to six “low-load” branches were tagged per tree, with the branches spaced throughout the canopy to normalize for sun exposure. One tagged inflorescence or nut cluster per tree was removed at each developmental time point studied and stored at 4 °C for further processing within 24 h. Trees bearing ‘Golden Hills’ scions were approximately 15 years old, while trees bearing ‘Kerman’ scions were approximately 20 years old. Both scion types were grafted onto the ‘Pioneer Gold I’ (*Pistacia integerrima*) rootstock. In 2022 in early April, fruit from six trees each of ‘Golden Hills’, ‘Lost Hills’, and ‘Kerman’ were sampled from an orchard located at Mendota, CA, USA, using the same methods as 2021. The trees were between 7 to 8 years old and grafted onto the ‘UCB1’ (*P. atlantica x P. integerrima*) rootstock.

In 2021 and 2022, randomly selected *P. vera* and *P. atlantica* trees from the United States Department of Agriculture (USDA) Germplasm collection at the Wolfskill Experimental Orchard (Winters, CA, USA) were used to validate the traits of interest. At the start of bloom in 2021, six *P. vera* trees were randomly selected, and six female inflorescences per tree were bagged prior to bud break. At the start of anthesis, each inflorescence was manually pollinated and enclosed in a wax paper bag for protection and to exclude extraneous airborne pollen. A subsampling of pollinated fruit was performed at 35 – 42 d post anthesis (dpa) for suture and cell morphology analyses, which corresponds to 200 – 640 growing degree days (GDD). In 2021, due to damage by pests, fauna, and an unexpected tree death, the final harvest was performed at 147 dpa (∼2298 GDD) instead of 154 dpa (∼2570 GDD), and only around half of the fruit clusters tagged were ultimately recovered from five of the six trees to use in analysis for kernel shape. In 2022, eight *P. atlantica* trees were randomly selected and fruit sampled at 35 – 42 dpa were used for suture anatomy and morphology analyses. The full list of GDD corresponding to the dpa and dates sampled are reported in Supplemental Table S1. *P. atlantica* blooms earlier than *P. vera,* and variations in climate between orchards and between years resulted in the variability in GDD and sampling date ranges presented. The monthly total precipitation, average air temperature, and average soil temperature from the three weather stations 7, 80, 105, (California Department of Water Resources 2023) closest to the commercial orchard sites sampled, as well as from Wolfskill Experimental Orchard (station 139), are reported for 2021 and 2022 in Supplemental Table S2. The accession information of the trees used is listed in Supplemental Table S3.

### Sample processing for microscopy

Fruit samples were excised into 5 mm thick sections in the transverse plane (Fig. 1) and fixed in 4% (w/v) paraformaldehyde (M26863; Thermo Fisher Scientific, Waltham, MA, USA) in phosphate buffered saline, 0.1% nonionic surfactant (Tween-20, J20605-AP; Thermo Fisher Scientific). Perfusion of the samples was achieved with a vacuum chamber (model 5530000; Labconco Corporation, Kansas City, KS, USA). Samples were incubated in fixative at a pressure of –90 to –100 KPa for 1.5 h at room temperature. For sectioning, the samples were trimmed down to a 3-5 mm region around the area of interest and embedded in 5% (w/v) agarose (A9539; Sigma-Aldrich, St. Louis, MO, USA) in deionized water. The embedded samples from 35 and 42 dpa were sectioned using a microtome with vibrating blade (Vibratome 1000 Plus Sectioning System; Technical Products International Incorporated, St. Louis, MO, USA) to 100 µm as previously described (Zhang S et al. 2021). A very limited number of 42 dpa sections with intact sutures were obtained using this method due to the lignified state of the endocarp at this stage, which rendered the shell less flexible and increased its tendency to break at the suture from the force of the blade.

### Assessment of suture and shell strength

The strength of the pistachio suture at each time point was measured at the longitudinal axis of fruit halved at the midpoint using the texture analyzer (TA.XT Plus Texture Analyzer; Texture Technologies Corporation, Hamilton, MA, USA) to exert compressive force against a metal platform. Fruit were halved in the frontal (coronal) plane at the nonsuture point, midway between the dorsal and ventral axis (Fig. 1). The kernel, when present, was removed before the pericarp was placed cut-face down on the instrument platform. The compression force (grams) required for breaking the pericarp along the suture was measured by compression using an aluminum cylindrical probe that is 2 inches in diameter (TA-25 probe; Texture Technologies Corporation) with pre-test speed of 2 mm·s^-1^ and test speed of 1 mm·s^-1^.

The strength of the pistachio shell at the nonsuture site was measured by placing the separated endocarp halves edge down on the platform. The force required for breaking the shell was measured as described above. The shells of fruit sampled immediately prior to commercial harvest in the orchard (133 dpa for ‘Golden Hills’, 154 dpa for ‘Kerman’, approximately 2422 and 2829 GDD, respectively, for 2021 at that field site) were used for the last timepoint measurement.

### Histological staining and microscopy

Calcofluor White stain (18909; Sigma-Aldrich) was used following the manufacturer’s instructions. The stain was mixed with 10% (w/v) potassium hydroxide in double deionized water in 1:1 ratio, and the samples were stained with this mixture for 1 min and rinsed once with deionized water before imaging. All fluorescence microscopy images were acquired using laser scanning microscopy (710 Axio Observer system; Zeiss, Oberkochen, Baden-Württemberg, Germany). A 408 nm laser (2% power) was used to excite the fluorophores, and Calcofluor White emission was collected over a wavelength range of 420 to 488 nm. Autofluorescence was captured using the same settings but by increasing the gain from 350-400 range to 600-700 range. Fluorescence images were obtained using three microscope objectives: the Plan-Apochromat 10x magnification 0.45 numerical aperture M27 thread size objective (420640-9900-000; Zeiss), the Plan-Apochromat 20x magnification 0.8 numerical aperture M27 thread size objective (420650-9902-000; Zeiss), or the LD C-Apochromat 40x magnification 1.1 numerical aperture M27 thread size objective (421867-9970-000; Zeiss). The pinhole was set at 1.88 Airy Units. The Zen 2011 SP3 (Zeiss) software was used for image acquisition and the Zen Lite software (ZEN 3.2, Zeiss) for image export.

### Shell sclerenchyma cell measurements

Maceration of shell was performed as described by Antreich et al. (2019). Briefly, 133 dpa fruit were cut in the transverse plane halfway between the apex and the base (pedicel end) and a 5 mm by 5 mm piece of endocarp was removed from the nonsuture site. The endocarp was submerged in 1:4:5 v/v/v mixture of 30% hydrogen peroxide (HX0635-2; EMD Millipore, Burlington, MA, USA), distilled water, and glacial acetic acid (A36S-212; Thermo Fisher Scientific) and heated at 60 °C for 72 h. The tissue was then rinsed three times with distilled water and stained with Calcofluor White for subsequent imaging with confocal microscope as described above.

### Fruit dimensions measurements

The fruit were sectioned in the transverse plane at the median (equidistant between the apex and base) of the fruit to expose the fruit median transverse section for imaging (Fig. 1). Fruit that contained no kernels, or “blanks,” were excluded from analyses. Measurements of suture angle, kernel shape, and shell thickness were performed manually using the “measure” tool in Fiji (Fiji Is Just) Image J (1.53f51) (Schindelin et al. 2012). For ease of discussion, “suture region” refers to the apical, dorsal, and ventral region, which is a broader region up to 500 μm x 500 μm in size that contains the suture. The term “suture” refers specifically to the suture itself and the three to seven layers of cells located at the suture along the interior-exterior axis, or the axis of dehiscence, of the pericarp.

### Data analysis and figure assembly

Data were recorded using either Microsoft Excel Office 365 (ver. 2305; Microsoft, Redmond, WA, USA) or Google Sheets (ver. 1.23.302.01.90; Google, Mountain View, CA, USA). Statistical analyses were performed using R (x 64 Windows version 4.0.3; R Core Team 2018) in RStudio (ver. 1.3.1093; RStudio Public Benefit Corporation, Boston, MA, USA) with the basic analysis of variance (ANOVA) and t-test functions. Emmeans (ver. 1.6.3) and multcomp (ver. 1.4-17) packages were used for least square means (LSM) post-hoc analyses (Hothorn et al. 2008; Lenth et al. 2023). Final graphs were generated using Microsoft Excel Office 365 and edited in image editing software Inkscape (ver. 1.0.1; Inkscape Project 2020). Figures were assembled in Inkscape.

## RESULTS

### Dorsal and ventral suture regions are asymmetrical in the transverse plane

Pistachio fruit shows bilateral symmetry along the longitudinal (apical-basal) axis. The fruit is flattened such that the two sutures running along the apical-basal axis are asymmetrical, with the dorsal suture showing greater curvature and, when sectioned in the transverse plane, a narrower (more acute) angle than the ventral suture (Fig. 1A). Earlier studies indicated that pistachio endocarp dehiscence initiates at the dorsal, but not the ventral, suture (Polito and Pinney 1999). To test the morphological and anatomical differences between the dorsal and ventral suture, we first determined the best sample orientation for our analyses. The endocarp dehisces along the suture and, when dehisced, splits the pericarp into two symmetrical halves (Fig. 1A). Therefore, in order to assay the difference caused by the asymmetry between the dorsal and ventral suture region, we focused our analyses on the transverse plane of section, such that the anatomy of the different tissue layers in each section can be clearly displayed (Fig. 1A, 1B). Micrographs throughout this study are presented with the exterior of the fruit in the upward direction (Fig. 1C). The term “suture” refers to the cells located at the suture, that undergo cell-cell separation, and the cells immediately adjacent, while the term “suture region” refers to the broader region of cells indicated in Fig. 1C.

### The dorsal suture shows a trend of weakening over time

Fruit dehiscence can be caused by a combination of mechanical force applied to the pericarp and the development of a mechanically weak region in the pericarp (Hofhuis et al. 2016; Ogawa et al. 2009; Parker et al. 2021). To determine whether mechanical weakness at the suture region contributes to the difference in the split rate (percentage of fruit with fully split shell) in ‘Golden Hills’ and ‘Kerman’, we assayed the suture strength of the two cultivars by measuring the compression force necessary to break the shell along the suture. We measured the dorsal suture, where dehiscence initiates, and compared it to the ventral suture. Both the dorsal and ventral suture region show a trend of decreasing suture strength during the later stages of fruit ripening, though there is no significant difference between ‘Golden Hills’ and ‘Kerman’ at the late stage immediately prior to harvest (Fig. 2A, 2B). However, when we compared dorsal strength at 133 dpa against 91 dpa, ‘Golden Hills’ showed a greater decrease in dorsal suture strength than ‘Kerman’ (*P* < 0.001 for dorsal, Two-way ANOVA, LSM α = 0.05, Fig. 2A). It is also worth noting that the suture strength at the dorsal region tends to be more variable than the ventral region, especially near the end of the developmental time points (Fig. 2A, 2B).

**Fig. 2.**
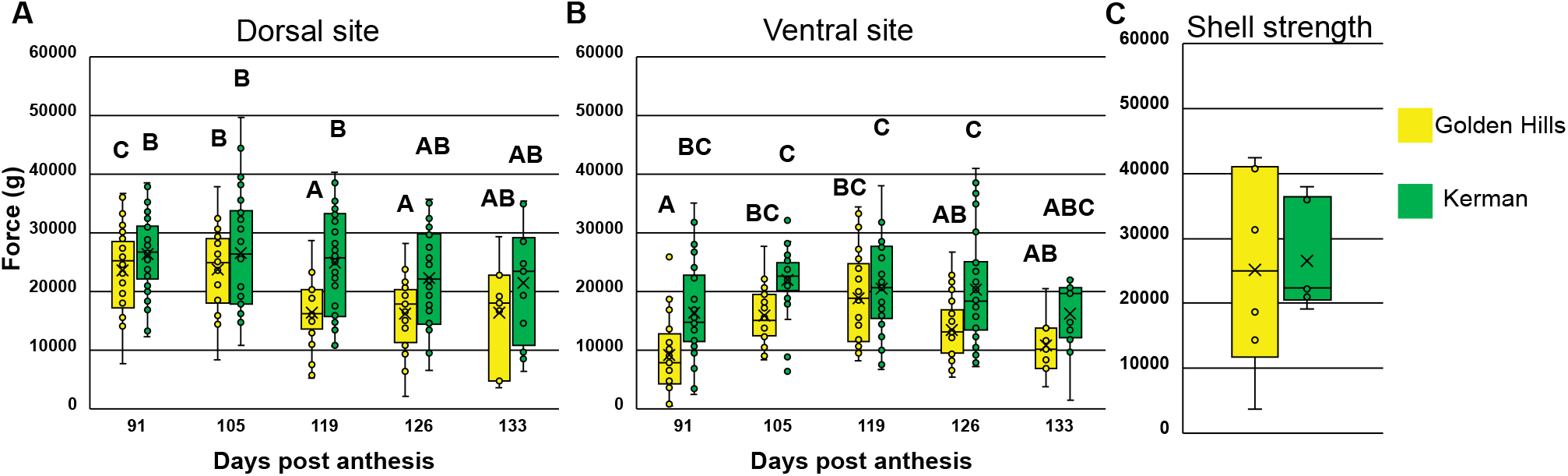
Pistachio shell strength measurement for ‘Golden Hills’ and ‘Kerman’ endocarp at dorsal suture (A), ventral suture (B), and nonsuture site (C) during late-stage fruit development. (A) Mechanical compressive force required to break dorsal suture in ‘Kerman’ and ‘Golden Hills’ fruit from 91 to 133 d post anthesis (dpa). *N* = 7-34 fruit from six trees per genotype. Probability not significant (NS) for interaction, ≤ 0.01 for genotype and dpa. Two-way analysis of variance (ANOVA). Different letters indicate significant differences under least square means (LSM) analysis with α cut off of 0.05. (B) Compressive Force required to break ventral suture of ‘Kerman’ and ‘Golden Hills’ fruit. *N* = 7-34 fruit from six trees per genotype from 91 to 133 dpa biweekly timepoints. Probability NS for interaction, < 0.01 for genotype and dpa. Two-way ANOVA. LSM α = 0.05. (C) Compressive force required for breaking endocarp of ‘Golden Hills’ and ‘Kerman’ fruit at nonsuture site at the final commercial harvest date, 133 dpa for ‘Golden Hills’ and 154 dpa for ‘Kerman’. Probability NS. *N* = 6 fruit from six trees. Two-tailed t-test.

### ‘Golden Hills’ and ‘Kerman’ show similar mechanical strength at nonsuture regions

The texture and flexibility of the endocarp at nonsuture sites may contribute to the stress experienced by the suture during endocarp dehiscence. There are many mechanical properties that can be investigated, but since it is ultimately the sum of all mechanical properties that contribute to endocarp split, rather than individually measuring parameters such as texture (“hardness”) or flexibility (“bendability”), we chose to directly measure the breakability of the two halves of the pistachio endocarp under pressure. To assess how much force is required for the endocarp to break at nonsuture areas, we measured the compression needed to break the two halves of the pistachio shell that were already separated along the suture. We found no significant difference between ‘Golden Hills’ and ‘Kerman’ at the date of final harvest (two-tailed t-test Fig. 2C), suggesting that the combined mechanical properties of the shell at nonsuture areas are similar in the two cultivars.

### Endocarp sclerenchyma cell size is smaller at the suture

Since size and shape of the sclerenchyma contribute to shell strength in other “nuts” (Huss et al. 2020), we investigated whether cells located at the suture of the pistachio endocarp are different from those at the suture adjacent region and nonsuture sites. Due to the mechanical strength of the highly lignified shell when compared to the relative fragility of the suture, we had difficulties obtaining thin sections of intact, unsplit sutures during late stages of development at the end of the harvest season. However, given that cell wall lignification is initiated by May, although shell hardness continues to increase afterwards (Polito and Pinney 1999; Shuraki and Sedgley 1996b; Zhang L et al. 2021), we reasoned that deposition of lignin indicates that the cells are at a stage where they are no longer expanding or undergoing dramatic changes in cell lobing. With the assumption that cell morphology is no longer changing in the shell once lignification has begun, we used sections of the endocarp at the relatively earlier stages of lignification to estimate the morphology of the cells during subsequent development. We analyzed sections of the endocarp at the suture from 35 – 42 dpa (last week of May / first week of June), representing the oldest stage with obtainable intact, lignified suture sections.

When we sampled the dorsal suture region of the endocarp at 35 – 42 dpa in the transverse plane (Fig. 3A), we observed that the cells of the dorsal suture were smaller and more flattened compared to the nonsuture cells in the suture regions, which we defined as suture adjacent cells. We found that the flattening of the suture cells appears to be oriented along the radial (interior-exterior) axis of the pericarp (Fig. 3B, 3C). This is of particular interest since in fruit that undergo pod shattering, long, lignified cells are oriented parallel to the axis of dehiscence (Parker et al. 2021). Therefore, to test whether there is significant flattening of endocarp cells at the suture and whether these flattened cells are aligned along the radial axis, we quantified two dimensions of the cell shape. The first dimension was “cell height”, which is the length of the cell along (parallel to) the radial axis (Fig. 3B, 3D). The second dimension was “cell width”, or length of the cell perpendicular to the radial axis, when viewed in the transverse plane (Fig. 3B, 3D). We found that there was significant flattening of the cells at the suture even compared to cells in the adjacent area only 100 μm from the suture. At the suture, the cells are shorter in the perpendicular to radial axis compared to parallel to radial axis by as much as 20 μm, leading to cells whose length in the perpendicular orientation is nearly half their length in the parallel orientation (*P* < 0.001 for major effect between suture and suture adjacent region, at both dorsal and ventral regions, two-way ANOVA, Fig. 3D-F). This leads to cells that are more elongated on the radial axis. When dehiscence occurs along the suture, the cells separate along the radial axis, running from the interior to the exterior of the endocarp, similar to the orientation of dehiscence zone fibers in arabidopsis (Dardick and Callahan 2014; Ogawa et al. 2009). This difference appears both at the dorsal and ventral suture. Furthermore, we observed that the flattening at the suture is significantly greater than at suture-adjacent areas (LSM analysis α = 0.05, Fig. 3C-F). Together, this indicates that while the endocarp is primarily composed of only one cell type, there is a modification of cell shape at the suture, where the cells are significantly smaller and flatter than those away from the suture.

**Fig. 3.**
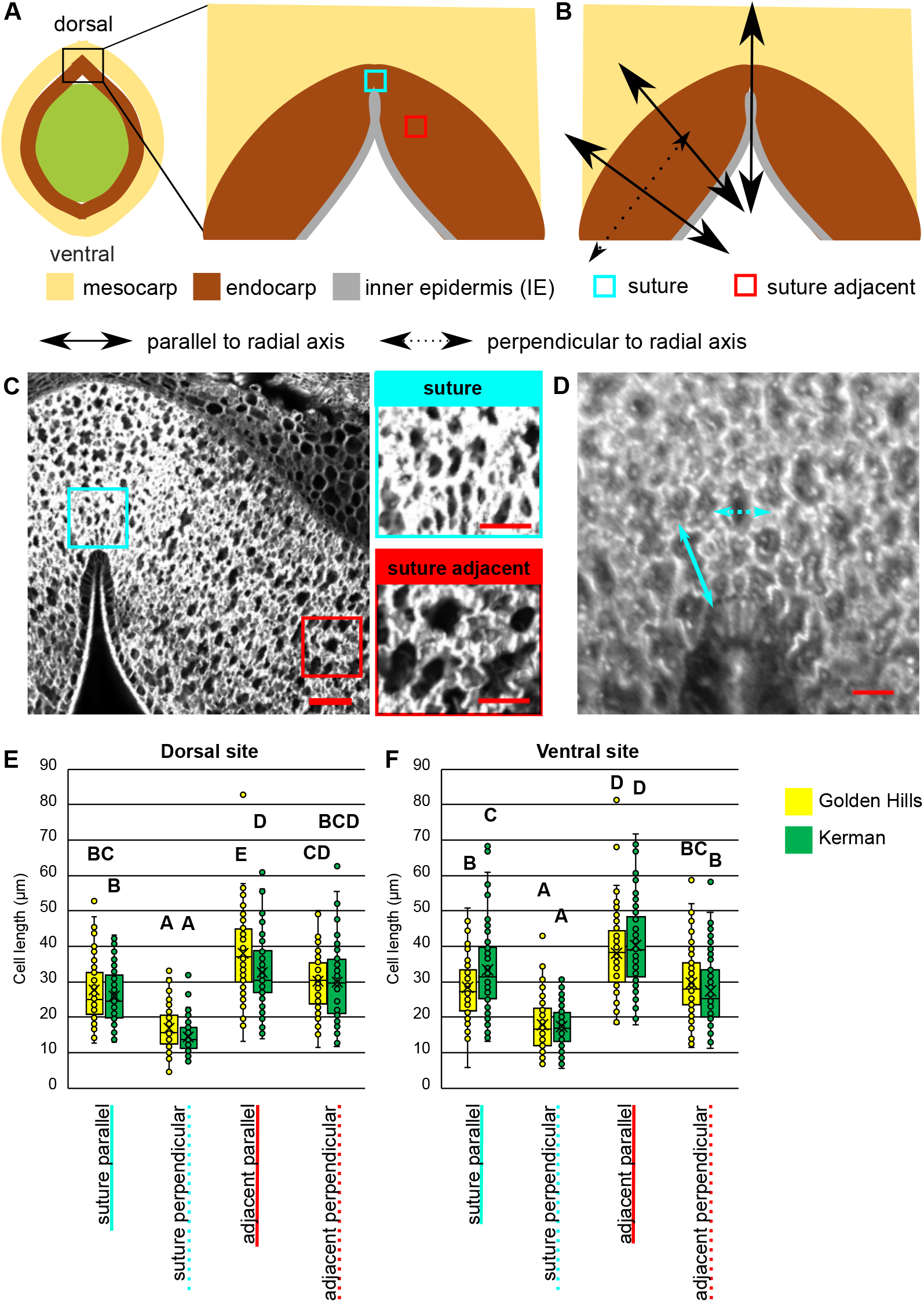
Anatomical analyses of pistachio endocarp sclerenchyma cell dimensions at suture and suture adjacent sites in ‘Golden Hills’ and ‘Kerman’. (A) Cells quantified are sampled from sections at the suture (cyan box) and suture adjacent (red box) regions 35-42 d post anthesis (dpa). (B) Two parameters, parallel to radial axis (dotted line) and perpendicular to radial axis (solid line), are measured for each cell. (C) Autofluorescence imaging of endocarp transverse section, located at suture at 35-42 dpa, excited using a 408 nm laser. Rectangles indicate sample regions corresponding to locations indicated in panel A. Cells appear more flattened at the suture (cyan square inset) compared to the suture adjacent site (red square inset) which appear wider with multiple lobes. Square insets of area are provided to show the representative cell morphology of each area. Scale bar= 100 μm for lower magnification, 50 μm for higher in insets. (D) Example of the lengths measured for each parameter of the cell, corresponding to axes indicated in the micrographs in (B). Scale bar = 20 μm. (E-F) Length measurements of endocarp cells at suture and suture-adjacent sites in 35 and 42 dpa ‘Golden Hills’ and ‘Kerman’ fruit. *N* = 51-60 cells per site from fruit of 4-5 trees per genotype. ** *P* < 0.01 between perpendicular and parallel to radial axis in suture and adjacent sites at apical, dorsal, and ventral locations, indicating differences in cell shape, two-way analysis of variance. Different letters indicate statistical difference in least square means analysis with an α of 0.05.

### Endocarp sclerenchyma cells are similar in ‘Golden Hills’ and ‘Kerman’

To further determine whether there may be other mechanical differences of the shell between ‘Golden Hills’ and ‘Kerman’ that were not detectable in the shell strength assays, we decided to directly assess the shape of the cells in the nonsuture region. Since it was difficult to capture the complex three-dimensional lobed shape in sections where all the cells are locked together, we macerated the shell tissue as earlier described by Antreich et al. (2019) in order to visualize individual cells. We then characterized the morphology of these macerated cells from nonsuture regions to determine whether lobing might contribute to split rate. We did not detect a significant difference in our microscopy-based assessment of cell size, lobe number, or lobe depth between ‘Golden Hills’ and ‘Kerman’ in endocarp sclerenchyma cells taken from mature fruit at 133 and 154 dpa, suggesting that there is no significant difference between the two cultivars in the morphology of cells that make up the bulk of the endocarp.

### Endocarp thickness at the suture may contribute to split rate

In our microscopy-based analysis of cell shape at the suture, we observed that there may be a decrease in endocarp thickness at the suture undetectable by the naked eye. Higher resolution micrographs allow for a better assessment of the endocarp and show that the endocarp can fold until the two sides are nearly touching at the dorsal suture (Fig. 4A, 4B). In micrographs of endocarp sections, we observed a significant difference in endocarp thickness between the suture and suture adjacent regions at both the dorsal and ventral sutures (two-way ANOVA *P* < 0.001 between site, < 0.01 between cultivar, LSM α = 0.05 between suture and suture adjacent areas, no significance found between cultivars, Fig. 4C, 4D). In addition, we observed that at the ventral suture, ‘Kerman’ endocarp was up to 200 μm (0.2 mm) thicker than ‘Golden Hills’ and this trend was observed both at the suture and suture adjacent region (*P* < 0.001 between site, two-way ANOVA, LSM α = 0.05, Fig. 4D). Together, these results show that both ‘Golden Hills’ and ‘Kerman’ have a thinner endocarp at the suture when compared to the suture adjacent area.

**Fig. 4.**
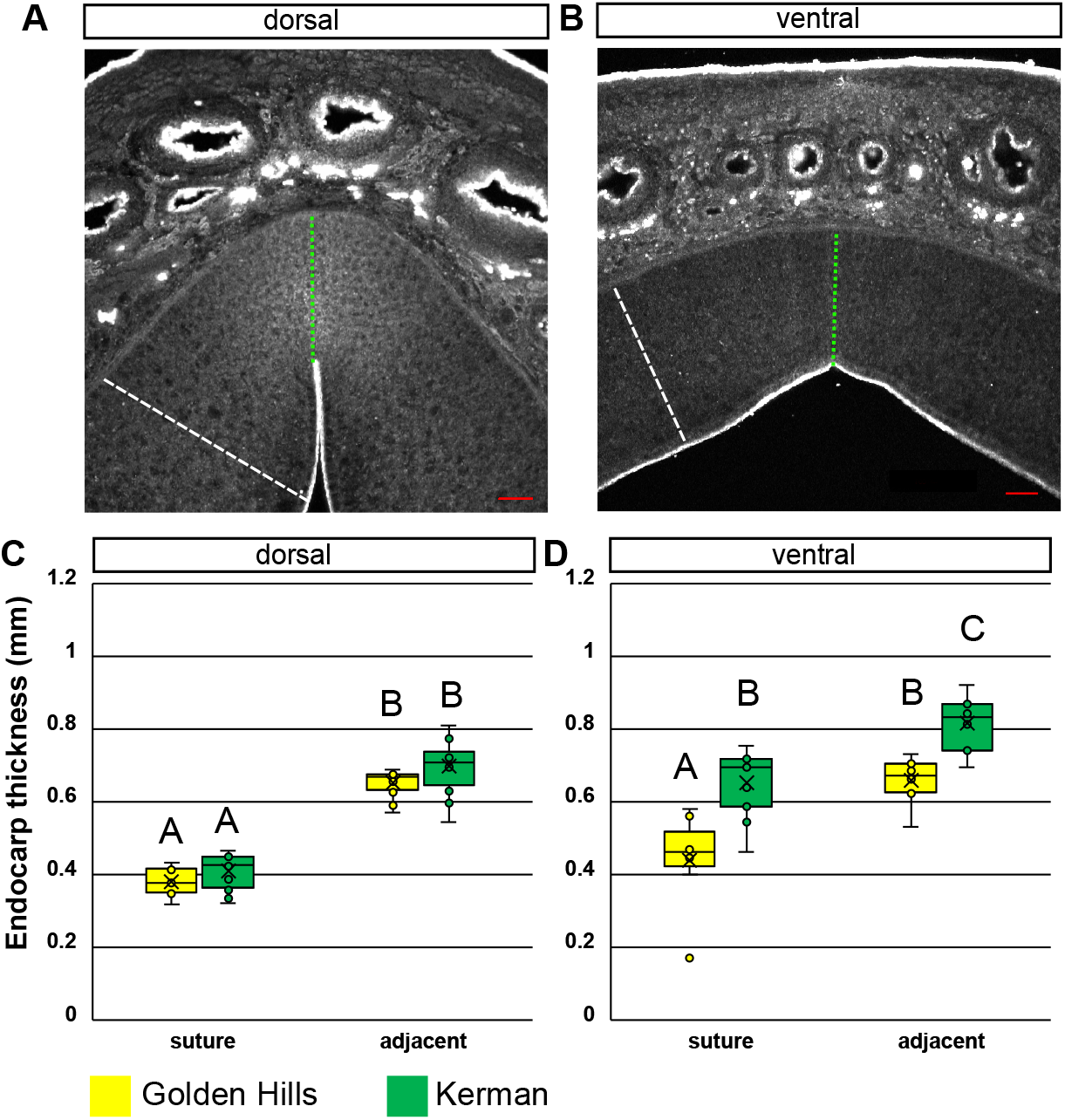
Anatomy and quantification of endocarp thickness at the dorsal and ventral suture in ‘Golden Hills’ and ‘Kerman’ fruit. Micrographs of autofluorescent pistachio endocarp, excited using a 408 nm laser, indicating the different locations measured for the dorsal (A) and ventral (B) suture and suture adjacent sites at 35-42 d post anthesis (dpa). Green dotted line indicates suture thickness, white dashed line indicates suture adjacent location thickness. Scale bars = 100 μm. (C, D) Quantification of endocarp thickness at dorsal and ventral sutures and suture adjacent sites from 35-42 dpa. *N* = 11-15 sections from four to five fruit. *P* < 0.01 for genotype and (suture versus adjacent) location at dorsal and ventral suture, no significance found for interaction, two-way analysis of variance. Different letters indicate statistical difference in least square means analysis with an α of 0.05.

Since a thinner endocarp at the suture region is implicated in endocarp dehiscence in pistachio (Shuraki and Sedgley 1996a), we measured the thickness of the suture and nonsuture regions in ‘Golden Hills’ (higher shell split rate) and ‘Kerman’ (lower shell split rate) to determine whether these traits are associated with the rate of endocarp dehiscence. Analysis of endocarp thickness at the suture and nonsuture regions using photographs of fruit sections showed a difference between cultivars at 133 dpa (*P* < 0.05 between genotype, two-way ANOVA, Supplemental Fig. 1). However, LSM analysis did not detect any significant difference between any of the groups (LSM analysis α = 0.05), likely due to the limits of the assay to detect such differences.

### Kernel width in transverse section is associated with increased endocarp dehiscence

Thinner endocarp and flattened cells at the suture may create mechanical weakness that facilitate shell split; however, the mechanical forces that can act on this weakness to cause dehiscence is unknown. Previous work by Polito and Pinney observed that “blank” fruit, which do not contain a kernel, did not dehisce, and posed the hypothesis that kernel expansion may provide the force behind endocarp dehiscence (Polito and Pinney 1999). If this is the case, kernel size or its dimensions should correlate with the rate of endocarp dehiscence. Therefore, we asked whether ‘Golden Hills’ had a larger kernel and a wider “girth” in the transverse plane when compared to ‘Kerman’. We observed in transverse sections cut at the median of the fruit that ‘Golden Hills’ had a “wider” kernel shape when compared to ‘Kerman’ (Fig. 5A).

**Fig. 5.**
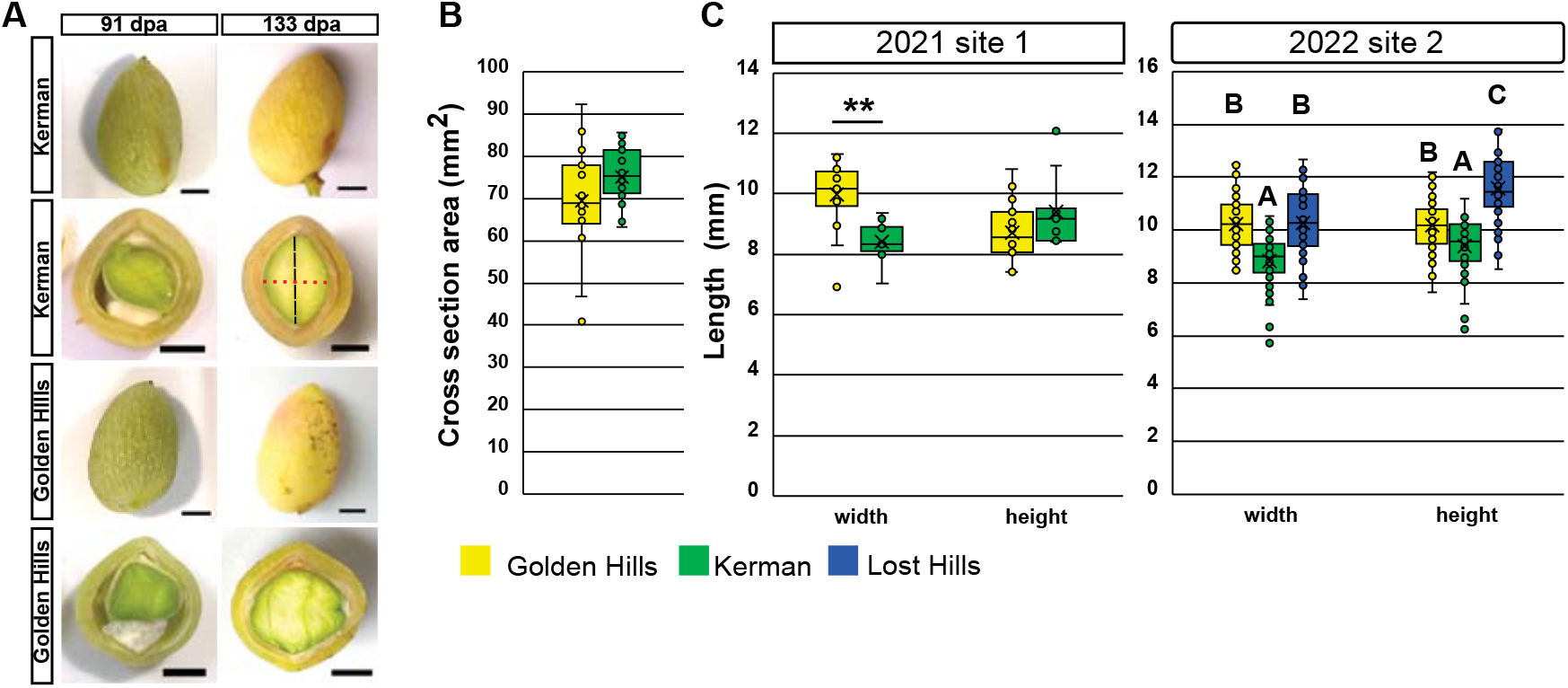
Pistachio kernel shape and dimensions measured in ‘Golden Hills’, ‘Lost Hills’ and ‘Kerman’ at median transverse section at late-stage fruit development. (A) Photographs of longitudinal and transverse section of ‘Golden Hills’ and ‘Kerman’ fruit from 91 to 133 d post anthesis (dpa). Red dotted line indicates the length measured for kernel “width”. Black dashed line indicates the length measured for kernel “height”, in transverse sections. Scale bars = 5 mm. (B) Average height and width of the kernel at the transverse section cut along the median of the fruit in ‘Golden Hills’, ‘Kerman’’, and ‘Lost Hills’ at 133 dpa for consecutive years in two sites. *P* < 0.01 for width, no significance (NS) for height for commercial orchard sampled in 2021 (Cantua Creek, CA, USA). *N* = 12-18 fruit from 6 trees per genotype. Two-tailed t-test. *P* < 0.01 for width and height for commercial orchard sampled in 2022 (Mendota, CA, USA). *N* = 38-56 fruit from six trees per genotype. One-way analysis of variance, different letters indicate significant differences with least square means analysis with α = 0.05. Data indicate consistent significant difference in width between genotypes (C) Average surface area of kernel cut in transverse section at median of the fruit at 133 dpa. Probability NS. *N* = 12-18 fruit from six trees per genotype, two-tailed t-test.

We then quantified kernel dimensions using the kernel transverse section area taken at the median of the fruit, halfway between the tip (the apex) and the base (the pedicel end). Since the pistachio kernel has a complex asymmetrical shape which varies even within a single tree, we reasoned that assuming the kernel fills the entire interior of the pericarp, selecting the median point between the tip and the base of the fruit would allow us to measure the kernel dimensions consistently between fruit and make comparisons at anatomically similar regions (Fig. 1A, 1B), even if the exact curvature of the kernel may vary between fruit. We found that the “width” (the sagittal axis) of ‘Golden Hills’ kernel transverse sections were 20% wider compared to ‘Kerman’ (*P* < 0.001, two-tailed t-test, Fig. 5B). There was no significant difference in the kernel “height” in the transverse plane (the dorsal-ventral axis) (Fig. 5B, two-tailed t-test). In other words, the major axis frequently lie on the sagittal axis for ‘Golden Hills’, while in ‘Kerman’ the major axis may more often lie in the coronal (frontal) axis when the fruit is viewed in the transverse plane. Interestingly, we did not find a significant difference in transverse sectional surface area between the two cultivars (Fig. 5C, two-tailed t-test), suggesting that the kernels did not seem to be significantly different in their girth in our plane of section.

‘Kerman’ has a longer growing season than ‘Golden Hills’ and is harvested later. During the commercial harvest of 2021, we sampled ‘Golden Hills’ at 133 dpa and ‘Kerman’ at both 133 and 154 dpa due to the difference in their commercial harvest dates. Thus, a concern was raised that the difference in width between the cultivars at 133 dpa is an artifact of the longer growing season of ‘Kerman’. To address this issue, in order to account for kernel growth in ‘Kerman’ post 133 dpa, we sampled and measured the ‘Kerman’ kernels again at 154 dpa (Supplemental Fig. 2A). We observed while ‘Kerman’ kernel width does increase between 133 and 154 dpa (*P* < 0.01, two-tailed t-test, Supplemental Fig. 2B), the kernel width of ‘Golden Hills’ at 133 dpa was still wider than that of ‘Kerman’ at 154 dpa (*P* < 0.01, two-tailed t-test, Supplemental Fig. 2C). In summary, ‘Golden Hills’ has a wider kernel shape compared to ‘Kerman’ at the final date of commercial harvest, which could contribute to endocarp dehiscence by serving as an expansion force that acts upon the mechanically weak points of the endocarp.

To further test our hypothesis, we expanded our analysis to an additional growing season in 2022 and included ‘Lost Hills’, which has a relatively high split rate similar to that of ‘Golden Hills’ (Parfitt et al. 2008). As expected, at the transverse section, the kernel width, but not height, correlated with split rate, with both ‘Lost Hills’ and ‘Golden Hills’ having wider kernels than ‘Kerman’ (*P* < 0.001, one-way ANOVA, Fig. 5B). In contrast, the height of the kernel was not strongly correlated with the split rate, with ‘Lost Hills’ showing a greater height than ‘Golden Hills’, and ‘Golden Hills’ showing a greater height than ‘Kerman’ (*P* < 0.001, one-way ANOVA, Fig. 5B).

### The pistachio endocarp is sharply folded at sutures where dehiscence initiates

The angle at the suture has been previously implicated in pistachio endocarp split rate, though hairpin-like folding at the dorsal suture, which forms a furrow-like shape, had not been quantified (Shuraki and Sedgley 1996a). Due to this furrowing, the dorsal suture has two angles: the furrow angle, or the acute folding that is only visible through the microscope, and the suture site angle, which is the suture ridge angle that is visible to the naked eye (Fig. 6A). It is unknown whether the furrowing or the suture site angle is associated with endocarp dehiscence.

**Fig. 6.**
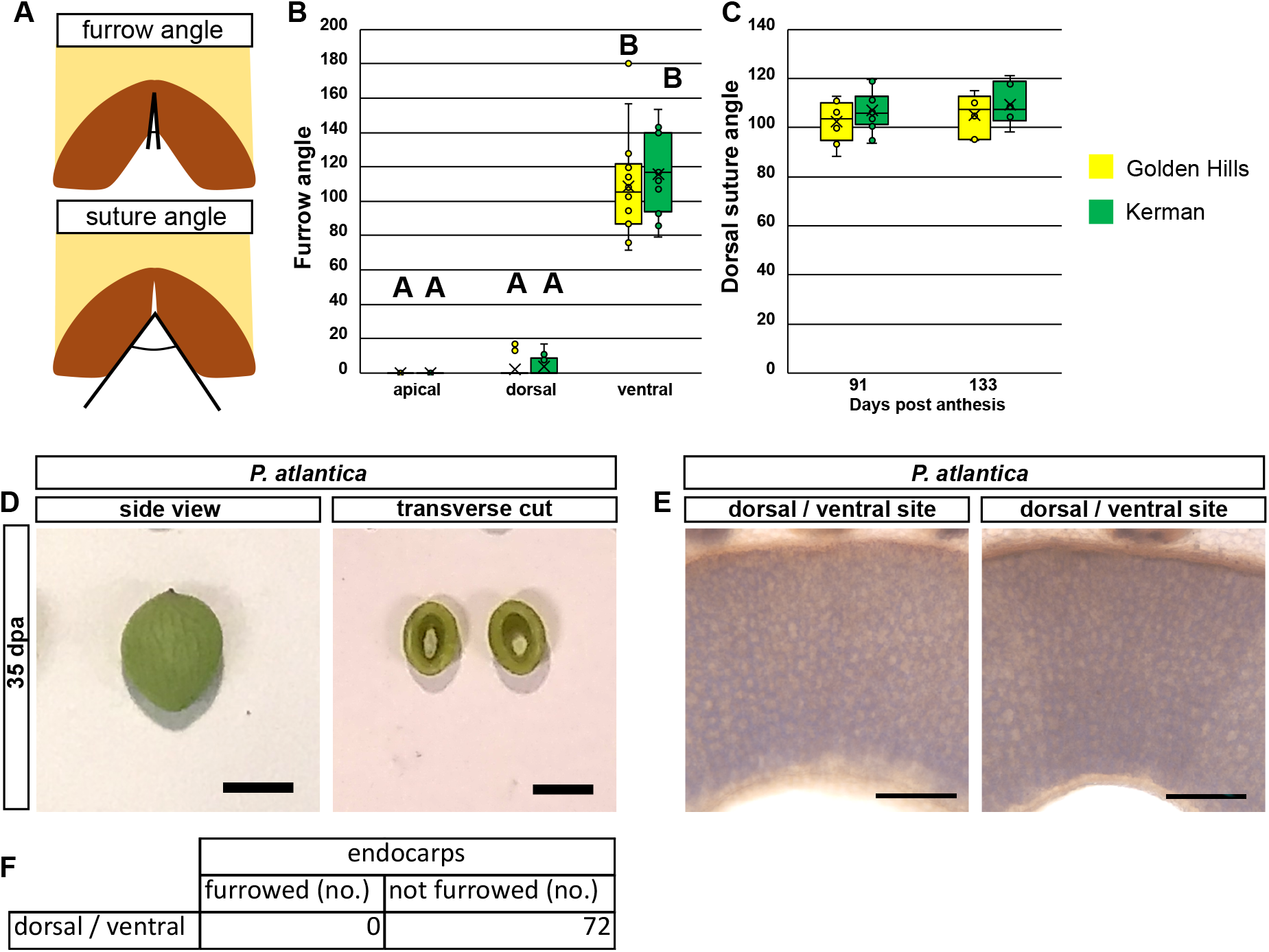
Anatomical analyses of dorsal and ventral suture angle in endocarp of *Pistacia vera* and *P. atlantica* at 35-42 d post anthesis (dpa). (A) Diagrams indicating the location of the furrow angle (top) and suture site angle (bottom) quantified in (B) and (C) for *P. vera* fruit. (B) Furrow angle at the apical, dorsal, and ventral sutures at 35-42 dpa in *P. vera* ‘Golden Hills’ and ‘Kerman’. *N* = 12-18 fruit per genotype for B, C, E, two-tailed t-test. *N* = 4-8 for apical, 11-13 for dorsal and ventral site per genotype for F, two-way analysis of variance, different letters indicate significant differences with least square means (LSM) analysis with α = 0.05. (C) Dorsal site angle of ‘Golden Hills’ and ‘Kerman’ at start and end of late-stage fruit development, 91-133 dpa. Probability not significant. (D) Representative photographs of the 35 dpa indehiscent *P. atlantica* fruit in side view and in transverse section cut at the median of the fruit. Scale bars = 5 mm. (E) Transverse sections of the two sutures of the *P. atlantica* at 35 dpa, stained with Evan’s Blue. Dorsal and ventral sites are virtually indistinguishable with no detectable acute-angle furrowing. Scale bars = 200 μm. (F) Furrowing is not observed in *P. atlantica* fruit. *N* = 48 suture sections from 36 fruit collected from eight trees.

Due to the difficulty of obtaining endocarp sections with an intact suture during the late stages of fruit development, we sampled endocarps from fruit between 35 and 42 dpa to examine the hairpin-like furrow angle (Figs. 4A, 6A). To test whether the furrow angle contributes to the difference in dehiscence between ‘Golden Hills’ and ‘Kerman’, we measured the dorsal, ventral, and apical furrow angle in micrographs. We observed that at the dorsal and apical suture, where dehiscence initiates, the furrows were often folded so that the inner epidermis of the two halves of the pistachio shell were pressed against and parallel to each other, forming a hairpin-like shape with a furrow angle of essentially zero (Table 1, Fig. 4A). In contrast, at the ventral suture, where dehiscence does not initiate, the two halves of the shell tend to form obtuse furrow angles (Table 1, Fig. 4B). This difference in occurrence of furrowing is significantly different between dorsal, apical, and ventral sutures (*P* < 0.001, Fisher’s exact test, Table 1). Quantification of these furrows showed that the ventral suture angle was significantly wider compared to apical and dorsal suture angles (*P* < 0.001, two-way ANOVA, Fig. 6B). There is no significant difference in the furrow suture angle between ‘Golden Hills’ and ‘Kerman’.

**Table 1.** Frequency of pistachio endocarp with acute-angle furrowing observed in transverse sections of fruits at apical, dorsal, and ventral suture. ‘Golden Hills’ and ‘Kerman’ fruit were sampled at 35-42 d post anthesis. Furrowing with acute angle is only observed at apical and dorsal suture sites. *P* < 0.01, Fisher’s exact test. *N* = 11-12 fruit sampled from five to six trees.

|  |  | Endocarps (no.) |  |
| --- | --- | --- | --- |
|  |  | Furrowed (no.) | Not furrowed (no.) |
| Location | Apical suture | 10 | 2 |
|  | Dorsal suture | 25 | 0 |
|  | Ventral suture | 0 | 23 |

We also tested whether the cultivar with the wider kernel has a wider dorsal suture site angle by measuring the suture site angle that is visible with the naked eye (Fig. 6C). We found no difference in the angle of the dorsal suture site between ‘Golden Hills’ and ‘Kerman’ at either 91 or 133 dpa (Fig. 6C, two-tailed t-test).

To verify the relationship of the morphological asymmetry between the dorsal and ventral suture, furrowing, and endocarp dehiscence, we next examined the fruit of *P. atlantica,* a rootstock cultivar for which the endocarp does not dehisce upon maturity. Our data showed that there was no difference in the suture morphology between the dorsal and ventral regions and no hairpin-like furrowing at either suture (Fig. 6D-F). In summary, these data demonstrates that there is furrowing at the apical and dorsal sutures, the sites of dehiscence initiation in the *P. vera* species, but not at the ventral suture. In contrast, the indehiscent fruit of *P. atlantica* did not display endocarp furrowing.

### Kernel dimensions in a genetically diverse population

To test whether the aforementioned traits are consistently associated with endocarp dehiscence in other cultivars, in 2021 we manually pollinated and harvested fruit from 5 genetically diverse *P. vera* trees located in the USDA germplasm collection at Wolfskill Experimental Orchard. We measured kernel width, cell morphology, furrow angle, and suture thickness, all of which we have observed to be associated to some degree with endocarp dehiscence or increased shell split rate. For assessment of kernel width, we compared all the fruit with split shells to all the fruit with unsplit shells. We measured the width and height of the kernel as previously described and found that the kernels of the fruit with split shells, or dehisced endocarps, are significantly wider than the unsplit fruit by approximately 15% to 18% (*P* < 0.01, two-tailed t-test, Supplemental Fig 3B). The height of the dorsal-ventral axis of the kernel was not significantly different between the split and unsplit fruit (Supplemental Fig. 3, two-tailed t-test).

To verify additional traits, we selected trees that had a dramatic difference in split rate, at 85.71% and 0%, respectively (tree 1 and 3, Supplemental Fig. 3A). With the addition of a tree with a middle range split rate of 44%, we designated these trees as high, medium, or low split trees with significant differences in split rate (*P* < 0.001, Fisher’s exact test, Supplemental Fig. 3A). When we assayed the cell morphology at the suture of fruit from these representative trees, we found that the sclerenchyma cells are indeed significantly smaller at both the dorsal and ventral suture. Furthermore, higher split rate was associated with smaller cells at the suture, especially at the dorsal suture (*P* < 0.001 between suture and suture adjacent location for both dorsal and ventral, *P* < 0.001 between genotype, interaction effect <0.001 at dorsal, two-way ANOVA, Supplemental Fig. 4A, 4B). Furrow angle and endocarp thickness followed the same pattern, where a more acute furrow angle and a thinner suture were correlated with an increase in shell split rate (*P* < 0.001 between genotypes for suture angle, *P* < 0.01 between genotype for suture thickness, two-way ANOVA, Supplemental Fig. 4C-E). Overall, these data confirm that the presence of a furrow with an acute angle and compact cells flattened along the radial axis contribute to the rate of shell split.

## DISCUSSION

There is limited understanding on how pistachio can split at the suture despite the lack of an anatomically specialized dehiscence zone. Although kernel expansion and suture angle have been proposed in the past as traits associated with pistachio endocarp dehiscence (Polito and Pinney 1999), there has been very little research to experimentally test these hypotheses, especially on commercially grown cultivars (Polito and Pinney 1999; Shuraki and Sedgley 1996a). In our study, we used ‘Golden Hills’ and ‘Kerman’, two of the most popular commercial cultivars that have different split rates (Kallsen et al. 2009; Parfitt et al. 2007), to investigate the anatomical traits associated with shell split. Our study led to the development of a model for endocarp dehiscence in pistachio. We found that when the endocarp is comprised of a single cell type, difference in cell and suture shape can create a specialized zone of mechanical weakness. The expansion force from the growing kernel, which is dependent on the shape of the kernel, can then act upon this zone of weakness to induce cell-cell separation at the suture.

### The mechanical property of the endocarp is not the main factor in endocarp dehiscence

Mechanical properties of the nutshell, such as strength and rigidity, can depend on the overall shell shape and thickness as well as the composition of cell types and their cell walls and shape (Antreich et al. 2019; Koyuncu et al. 2004; Shuraki and Sedgley 1996a; Xiao et al. 2020; Zhao et al. 2019). The pistachio shell, or endocarp, is primarily composed of a type of single polylobate sclerenchyma cell, similar to what is observed in walnuts (Antreich et al. 2019; Fabbri et al. 1998; Polito and Pinney 1999; Shuraki and Sedgley 1996a; Xiao et al. 2020). Thus, the anatomical attributes of that single cell type, such as its shape and size, determine the mechanical properties of the shell. Our analysis of suture strength, shell strength, and the average endocarp cell morphology suggests that the mechanical properties of the endocarp as a whole are unlikely to be the main driver behind endocarp dehiscence (Fig. 2, Supplemental Fig. 5).

The consistently weaker suture of ‘Golden Hills’ (Fig. 2) and the gradual decrease in suture strength during ripening suggest that it is the mechanical properties of the suture, rather than the entire endocarp, that may contribute to endocarp dehiscence. Additional studies such as biochemical analyses of cell wall components at the suture site versus the nonsuture site are needed to determine whether the decrease in suture strength during 91-133 dpa involves mechanisms beyond changes in kernel expansion force. Modification in cell walls such as the breakdown of cellulose and pectin has been proposed as a mechanism to decrease the strength of dehiscence zones in multiple species (Clements and Atkins 2001; Roberts et al. 2002; Spence et al. 1996). To investigate cell wall modification at the suture, methods such as laser-microdissection of the cell layers immediately adjacent to the dehisced suture followed by biochemical analysis of the suture versus nonsuture region may provide more insights.

### Expansion force from kernel width may be associated with endocarp dehiscence at the suture

Premature rupture of carpel valves in the *fruitful (ful-1)* arabidopsis mutant has been attributed to the expansion force from crowded seeds (Gu et al. 1998). Similarly, based on the observation that pistachio nuts with no kernel do not show endocarp splitting, it has been proposed that kernel expansion may provide the mechanical force necessary for endocarp dehiscence (Polito and Pinney 1999). We observed that ‘Golden Hills’, which has a higher split rate in our study, has “wider” kernels than ‘Kerman’ and that fruit with split endocarps from the genetically diverse germplasm collection have wider kernels than fruit with unsplit endocarps (Figs. 5, 7). Furthermore, the difference in kernel width between ‘Golden Hills’ and ‘Kerman’ is roughly 10% to 15%, which is within the range of the 10% to 15% difference in split rate between the two cultivars (Kallsen et al. 2009, 2014).

The dimensions of the pistachio kernel can be defined by the height (dorsal-ventral axis), width (sagittal axis), and length (transverse, or apical-basal axis). Observations by L. Zhang et al. (2021) indicated that ‘Golden Hills’ and ‘Kerman’ kernels are similar in their lengths from the apex to the base of the fruit. This suggests that the expansion force from the kernel is unevenly distributed, with the force mainly concentrated in the sagittal axis. Therefore, the width of the kernel may be more critical to endocarp dehiscence than the height since it exerts a greater force perpendicular to the axis of dehiscence. Thus, although the width of the kernel is applying pressure directly to the nonsuture site, this could lead to tension build up at the suture, forcing the two halves of the endocarp apart. Although the greater kernel width and unchanged kernel height would predict an increase in transverse sectional surface area, we did not detect such a significant difference between ‘Golden Hills’ and ‘Kerman’ in transverse sections taken from the median of the fruit, halfway between the apical and basal ends (Fig. 5C). This may be due to the complex shape of the kernel, where the degree of asymmetry and curvature between the dorsal and ventral halves can create shapes where the largest transverse section area is located at different points along the apical-basal axis. This can potentially generate a difference in expansion force but without a difference in area at the median of the fruit (Fig. 8A).

The lack of difference in suture and furrow angles at the dorsal sites between ‘Golden Hills’ and ‘Kerman’, despite the change in the “width” of the kernel, suggests that expansion force may not be sufficient to distort the overall shape of the shell. The corollary to this is that the expansion force from the kernel may be stored as potential energy in the shell in the form of tension, similar to what is found in a compressed spring, and act as a source of stress on the mechanically weak region in the endocarp. Biophysical modeling, characterization of cell walls at the suture region, and tensile tests at the suture region will be necessary to separate the tension stored at the suture region from the mechanical weakness of the suture region.

### Cell and suture shape may contribute to form physically weak points at suture sites

In walnut, the lobed shape of the polylobate sclerenchyma cells is hypothesized to strengthen the shell by enabling cells to interlock tightly with each other, allowing the shell to remain intact on the macroscopic level even when microscopic cracks may appear along the middle lamella (Antreich et al. 2019; Xiao et al. 2020). When we quantified the shape of the cells in the pistachio endocarp, we observed that cells at the suture tend to be smaller compared to cells at the suture adjacent and nonsuture sites, and that the cells appear flattened in the radial plane. This is consistent in ‘Golden Hills’, ‘Kerman’, and the genetically diverse cultivars from the National Germplasm Repository-Davis, USDA germplasm collection (Davis, CA, USA) (Figs. 3, 7). Smaller cells may allow easier propagation of cracks (Huss et al. 2020). In addition, smaller cell size at the suture may also be the reason why the endocarp is thinner at the suture, which in turn exacerbates the region’s mechanical weakness (Fig. 4, Supplemental Fig. 3). The shape of lignified cells in dehiscence zones is important for fruit dehiscence in arabidopsis and legumes, where the fruit pericarp splits along the major axis (Dardick and Callahan 2014; Ogawa et al. 2009; Parker et al. 2020). Therefore, the flattening of the cells at the suture, which results in cells that are elongated along the interior-exterior axis of the fruit, may contribute to splitting at the suture along the radial axis.

In addition, furrowing of the endocarp at the apical and dorsal sites may be functionally similar to “fold and tear” lines on packaging and envelopes, providing yet another mechanism for creating a region of endocarp weakness. The angle of the suture has been previously implicated in shell split for fruit of *P. atlantica* and *P. vera* crosses (Shuraki and Sedgley 1996a). Indeed, the endocarp furrowing and the acuteness of the furrow at the dorsal suture consistently correlate with higher split rate in all the genotypes we studied (Fig. 6, Supplemental Fig. 3), suggesting that while kernel width may affect the split rate, the presence of the furrow may be important for initiation of endocarp split. The relationship between angle and strength has been studied at a ultrastructural level for the orientation of cellulose microfibrils, at the tissue level in *Cardamine* and *Bauhinia* species for lignified fiber orientation, and at the organ level in the walnut shell positions relative to the suture (Hofhuis et al. 2016; Koyuncu et al. 20024; Parker et al. 2021). Our data further cements this relationship by linking acute furrow angle to endocarp dehiscence. Optimal angles, such as the ones found in the hexagon of beehives, are known to occur in nature. The optimal angle for suture strength in pistachio is currently unknown. Since the kernel-derived mechanical force is applied from the interior of the angle, this may put stress on the “joint” where the “joint” is bent at an angle that is not physically optimal for bearing stress.

Thus, we propose a new mechanism for endocarp splitting in addition to the commonly known mechanism involving an anatomically distinct dehiscent zone defined by specialized cells and tissues. Namely, we suggest that a region of the pericarp can become a zone of dehiscence that is mechanically weak due to the thickness of the tissue and the size and shape of the cells (Fig. 7B). Future studies can address how much the cell size, cell flattening, and suture thickness each contribute to the overall strength at the suture. It is important to note that due to the shape of the pistachio fruit, the physical force at the dorsal and ventral sutures is unequally distributed along the entire curvature from the tip to the base of the fruit. Computational modeling may be required to examine how the uneven force exerted by the kernel is distributed along the length of the suture and how the strength of the suture affects overall endocarp dehiscence rate. For the low split-rate cultivars already planted in the field, studies on field management practices that can affect the rate of endocarp cell expansion and kernel expansion will be needed to assess how changes in endocarp sclerenchyma cell size and the timing and dimensions of kernel expansion may affect shell split-rate.

**Fig. 7.**
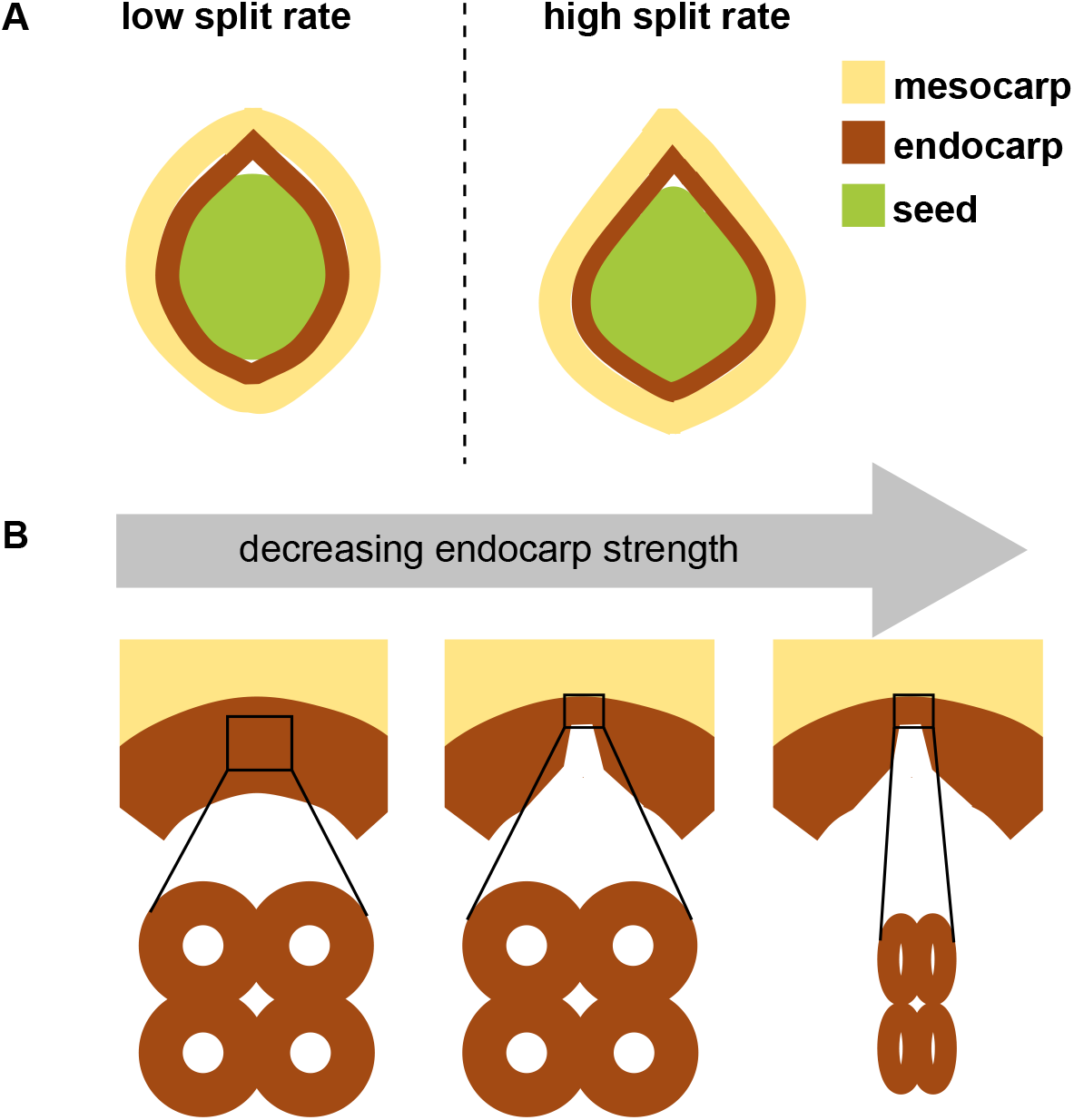
Model of effect of kernel shape and endocarp cell and suture morphology on pistachio shell split. The overall kernel shape in combination with suture morphology and suture cell shape can contribute to shell split. (A) Difference in expansion force generated by difference in kernel shape can contribute to split rate as defined by percentage of “full split” nuts between cultivars. (B) The presence of acute-angle furrowing at the suture, in combination with smaller, more flattened cells, creates a mechanically weak region in the endocarp, leading to a dehiscence zone which is specialized based on cell and shell shape rather than cell type.

To our knowledge, pistachio is the only “nut” that naturally dehisces upon maturity (Huss et al. 2020; Polito and Pinney 1999). Our study suggests that local variation in cell morphology, even when there is only one cell type present, may create mechanical weak points that, when acted on by the expansion force from the growing seed, can cause split in fruit pericarps.

