## supplemental table captions for "Difference in Kernel Shape and Endocarp Anatomy Promote Dehiscence in Pistachio Endocarp"

Supplemental table 1. Estimated growing degree days (GDD) for pistachio calculated based on temperature data corresponding to each day post anthesis (dpa) time point reported in this manuscript. Each dpa corresponds to a range in GDD due to differences in flowering time and phenology between the various cultivars in commercial orchards and the trees in the US Department of Agriculture germplasm collection in Wolfskill Experimental Orchard (Winters, CA, USA).

Supplemental table 2. Precipitation and temperature data from California Irrigation Management Information System weather stations for the 2021 and 2022 field sites. Stations 7, 80, and 105 are the stations closest to the commercial orchard sites in 2021 and 2022. Station 139 is located at Wolfskill Experimental Orchard (Winters, CA, USA).

Supplemental table 3. Inventory identification (ID), tree ID assigned in the manuscript, and location accession identification (location ID) of *Pistacia* trees sampled from US Department of Agriculture pistachio germplasm collection located at Wolfskill Experimental Orchard (Winters, CA, USA).
