## supplemental table 1 for "Difference in Kernel Shape and Endocarp Anatomy Promote Dehiscence in Pistachio Endocarp"

| dpa (d) | GDD | Date |
|---|---|---|
| 0 | 0 | 29 Mar - 14 Apr |
| 21 | 200-330 | 19 Apr - 5 May |
| 28 | 299-400 | 26 Apr - 12 May |
| 35 | 407-527 | 3 May - 19 May |
| 42 | 497-640 | 10 May - 26 May |
| 91 | 1500-1580 | 28 Jun - 14 Jul |
| 133 | 2280-2420 | 9 Aug - 25 Aug |
| 147 | ~2298 | 23 Aug - 16 Sep |
| 154 | ~2570 | 15-Sep |
