## supplemental table 2 for "Difference in Kernel Shape and Endocarp Anatomy Promote Dehiscence in Pistachio Endocarp"

| | Total precipitation (mm) | | | | Average air temperature (°C) | | | |
|---|---|---|---|---|---|---|---|---|
| Date | Station 7 | Station 80 | Station 105 | Station 139 | Station 7 | Station 80 | Station 105 | Station 139 |
| Jan 2021 | 70.8 | 105.2 | 65.8 | 79.4 | 8.8 | 8.9 | 8.6 | 9.4 |
| Feb 2021 | 2.1 | 10.6 | 0.7 | 12.7 | 11.3 | 10.7 | 11.2 | 12.2 |
| Mar 2021 | 28.7 | 44.5 | 14.7 | 20.5 | 12 | 11.8 | 11.8 | 12.4 |
| Apr 2021 | 3.3 | 5 | 0.7 | 3.8 | 16.9 | 17.3 | 17.4 | 17 |
| May 2021 | 1.8 | 0 | 0.1 | 0.2 | 21.1 | 21.2 | 22.2 | 21.2 |
| Jun 2021 | 0 | 0 | 0 | 0 | 25 | 26.2 | 26.5 | 23.5 |
| Jul 2021 | 0 | 0 | 0 | 0.2 | 27.3 | 29.5 | 29.6 | 23.8 |
| Aug 2021 | 0 | 0 | 0 | 10.4 | 25.5 | 27.5 | 27.6 | 24 |
| Sep 2021 | 0 | 0 | 0.3 | 1.5 | 23.4 | 24 | 24.6 | 22.4 |
| Oct 2021 | 35.6 | 36.7 | 14.6 | 152.6 | 16.9 | 16.2 | 17.2 | 15.6 |
| Nov 2021 | 5 | 14.7 | 3.6 | 26.4 | 12.3 | 12.2 | 12.1 | 12 |
| Dec 2021 | 37.2 | 108.3 | 39.1 | 165.4 | 7.9 | 8.1 | 7.7 | 7.6 |
| Jan 2022 | 0.2 | 0 | 1.5 | 9.3 | 8.7 | 7.9 | 8.1 | 8.8 |
| Feb 2022 | 20.1 | 2.7 | 0.9 | 0.7 | 10.5 | 9.2 | 10 | 11.2 |
| Mar 2022 | 35.1 | 23.1 | 10.6 | 13.1 | 14.5 | 14 | 14.5 | 14.3 |
| Apr 2022 | 10.6 | 11.1 | 5.8 | 13.7 | 16.4 | 16.2 | 16.9 | 16 |
| May 2022 | 0.1 | 0 | 0 | 4.1 | 20.2 | 20.8 | 21.2 | 20.3 |
| Jun 2022 | 0.7 | 0 | 0 | 14.7 | 25.2 | 26.2 | 26.5 | 24.6 |
| Jul 2022 | 0 | 0 | 0 | 17.1 | 26.6 | 28.1 | 28.5 | 24.4 |
| Aug 2022 | 0 | 3.5 | 0 | 20.2 | 26.6 | 28.5 | 28.8 | 24.8 |
| Sep 2022 | 1.1 | 5.6 | 6 | 32.5 | 25 | 25.8 | 26.3 | 23.5 |
| Oct 2022 | 0 | 0.5 | 0 | 6 | 19.1 | 19.1 | 19.9 | 18.1 |
| Nov 2022 | 19.3 | 19 | 0 | 23.1 | 9 | 8.8 | 8.9 | 9.4 |
| Dec 2022 | 74.8 | 136.3 | 46.6 | 172.1 | 7.5 | 7.8 | 7.7 | 7.1 |

| Average soil temperature (°C) | | | |
|---|---|---|---|
| Station 7 | Station 80 | Station 105 | Station 139 |
| 7.8 | 11.1 | 9.7 | 9.7 |
| 10.4 | 11.4 | 11.5 | 11.1 |
| 13.2 | 11.9 | 13.3 | 12 |
| 18.3 | 15.2 | 19.8 | 16.8 |
| 20.9 | 18.9 | 25.8 | 20.6 |
| 24.7 | 22.4 | 31.6 | 23.6 |
| 26.2 | 24.1 | 34.2 | 24.2 |
| 24.9 | 23.5 | 32.4 | 23.4 |
| 22.7 | 22.1 | 28.7 | 21.5 |
| 17.3 | 18.6 | 21.5 | 17.5 |
| 13.2 | 16.9 | 15.3 | 14.5 |
| 9.6 | 13.5 | 10.3 | 10.7 |
| 8.3 | 11.5 | 9.7 | 9.5 |
| 9 | 11 | 11.1 | 9.9 |
| 13.9 | 12.5 | 16.3 | 12.5 |
| 16.8 | 14.7 | 20 | 15 |
| 19.8 | 16.9 | 25 | 17.7 |
| 23.5 | 21.2 | 30.4 | 20.9 |
| 24.8 | 24.2 | 34 | 23.2 |
| 25.2 | 25.4 | 34 | 24 |
| 23.8 | 24.3 | 30.9 | 21.8 |
| 19.1 | 21.3 | 25.1 | 18.3 |
| 9.5 | 15.6 | 13.5 | 11.8 |
| 8.3 | 12.5 | 9.9 | 8.9 |
