## supplemental table 3 for "Difference in Kernel Shape and Endocarp Anatomy Promote Dehiscence in Pistachio Endocarp"

| Species used | Tree ID in manuscript | USDA Inventory ID | USDA location ID |
|---|---|---|---|
| *Pistacia vera* | 1 | DPIS 347.0000B | WEO D 07 24 |
| *Pistacia vera* | 2 | DPIS 245.0004A | WEO D 03 15 |
| *Pistacia vera* | 3 | DPIS 245.0003A | WEO D 03 14 |
| *Pistacia vera* | 4 | DPIS 261.0009A | WEO D 06 08 |
| *Pistacia vera* | 5 | DPIS 245.0006A | WEO D 03 17 |
| *Pistacia atlantica* | 1 | DPIS 196.0006A | WEO C 08 09 |
| *Pistacia atlantica* | 2 | DPIS 200.0009A | WEO C 08 19 |
| *Pistacia atlantica* | 3 | DPIS 200.0021A | WEO C 08 20 |
| *Pistacia atlantica* | 4 | DPIS 201.0005A | WEO C 08 30 |
| *Pistacia atlantica* | 5 | DPIS 201.0008A | WEO C 08 32 |
| *Pistacia atlantica* | 6 | DPIS 201.0012A | WEO C 08 35 |
| *Pistacia atlantica* | 7 | DPIS 350.0000A | WEO D 02 12 |
| *Pistacia atlantica* | 8 | DPIS 350.0000B | WEO D 08 19 |
