## supplemental figure captions for "Difference in Kernel Shape and Endocarp Anatomy Promote Dehiscence in Pistachio Endocarp"

Supplemental Fig. 1. Quantification of pistachio endocarp thickness in ‘Golden Hills’ and ‘Kerman’ at dorsal suture and nonsuture sites from photographs of fruit sections at 133 d post anthesis (dpa). (A) Transverse section of ‘Kerman’ fruit at 133 dpa at median of the fruit. Red lines indicate length measured for suture and nonsuture region. Scale bar = 5 mm. (B) Endocarp thickness at dorsal suture site and nonsuture site at 133 dpa. $P < 0.05$ for genotype at 133 dpa, no significance found for site and interaction. $N$ = 8-12 fruit from six trees per genotype. Two-way analysis of variance.

Supplemental Fig. 2. Quantification and comparisons of pistachio kernel dimensions at final date of commercial harvest. (A) ‘Kerman’ kernel width and height 154 d post anthesis (dpa). $N$ = 11 fruit from six trees. (B) Increase in ‘Kerman’ kernel width from 133dpa to 154 dpa. $P < 0.01$, two-tailed t-test. $N$ = 11 fruit from six trees. (C) 133 dpa ‘Golden Hills’ kernel width compared against 154 dpa ‘Kerman’ kernel width. $P < 0.01$, two-tailed t-test. $N$ = 11 ‘Kerman’, 18 ‘Golden Hills’. The kernel width of ‘Golden Hills’ at 133 dpa was wider than that of ‘Kerman’ at 154 dpa.

Supplemental Fig. 3. Quantification of number of pistachio fruit with split and not split shell found in fruit harvested from each tree and the tree’s shell split rate (A) and kernel height and width in transverse section of the fruit (B) from fruit of five trees in the US Department of Agriculture germplasm collection at Wolfskill Experimental Orchard (Winters, CA, USA) in 2021 harvested at 147 d post anthesis. $N$ = 17 for split shell fruit, 27 for unsplit shell fruit, from five trees. $P < 0.01$, two-tailed t-test.

Supplemental Fig. 4. Anatomical quantifications of pistachio shell sclerenchyma cells (A, B), suture furrow (C), and endocarp thickness (D, E) at the dorsal and ventral suture from fruit collected from the

US Department of Agriculture germplasm collection at Wolfskill Experimental Orchard (Winters, CA, USA). Measurements were taken from transverse sections of 35 d post anthesis fruit, at the median of the fruit, from 3 trees with high (85.7%), medium (44%), and low (0%) split rate. Lengths of the cells are measured perpendicular and parallel to the radial axis at the suture and suture-adjacent sites for sclerenchyma cell quantifications (A, B). $N = 50$ cells per site from five to six fruit from each tree type for A, B. $N = 5\text{-}6$ fruit for each tree type for C-E. For A, C, D $P < 0.01$ for comparison between trees, between sites, and interaction. For B, $P < 0.01$ for trees, sites, no significance (NS) is found for interaction. For E, $P < 0.01$ for trees, $< 0.05$ for sites, NS for interaction. Two-way analysis of variance. Different letters indicate significant differences with least square means analysis with $\alpha = 0.05$. More acute furrow angle and smaller cells at the suture are associated with higher endocarp split rate.

Supplemental Fig. 5. Quantification of pistachio endocarp sclerenchyma cell dimensions in ‘Golden Hills’ and ‘Kerman’ with focus on lobe characteristics. (A) Micrographs of calcofluor stained sclerenchyma cells of ‘Golden Hills’ and ‘Kerman’ whole endocarp maceration at date of final harvest (133 and 154 d post anthesis). Red dotted line indicates length of measurement for major lobe depth. Scale bars = 50 µm. (B-D) Quantification of macerated cells represented in (A). No significant difference was detected in major lobe depth, major lobe number, and average cell area between ‘Golden Hills’ and ‘Kerman’. $N = 23\text{-}34$ cells from six fruit sampled from six trees per genotype. Two-tailed t-test.
