## Supplementary figures and images for "Difference in Kernel Shape and Endocarp Anatomy Promote Dehiscence in Pistachio Endocarp"

### supplemental figure 1

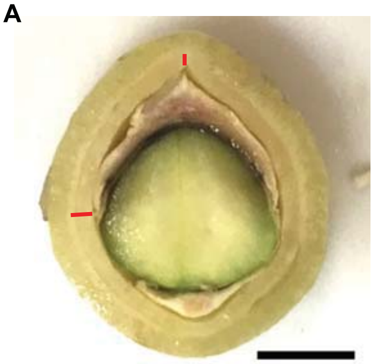

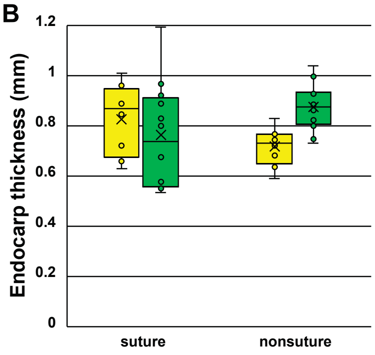

### supplemental figure 2

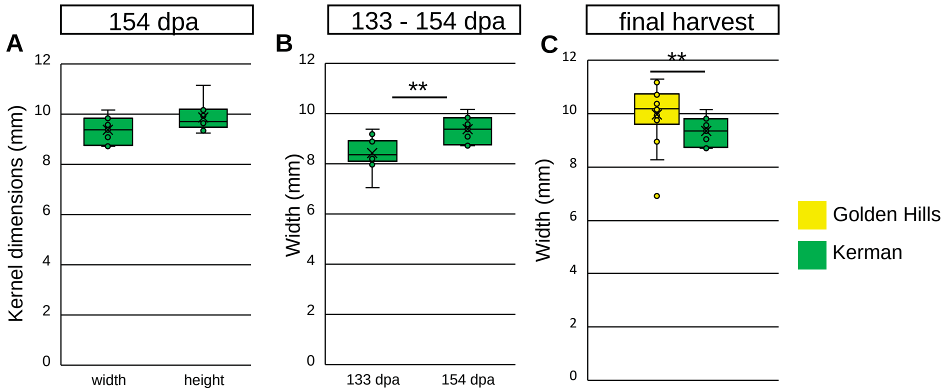

### supplemental figure 3

**A**

| tree ID no. | split (no.) | not split (no.) | split rate (%) |
|---|---|---|---|
| 1 | 18 | 3 | 85.71 |
| 2 | 11 | 14 | 44 |
| 3 | 0 | 12 | 0 |
| 4 | 0 | 1 | 0 |
| 5 | 11 | 6 | 64.71 |

**B**

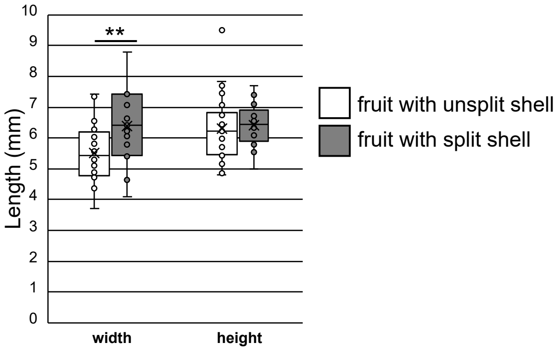

### supplemental figure 4

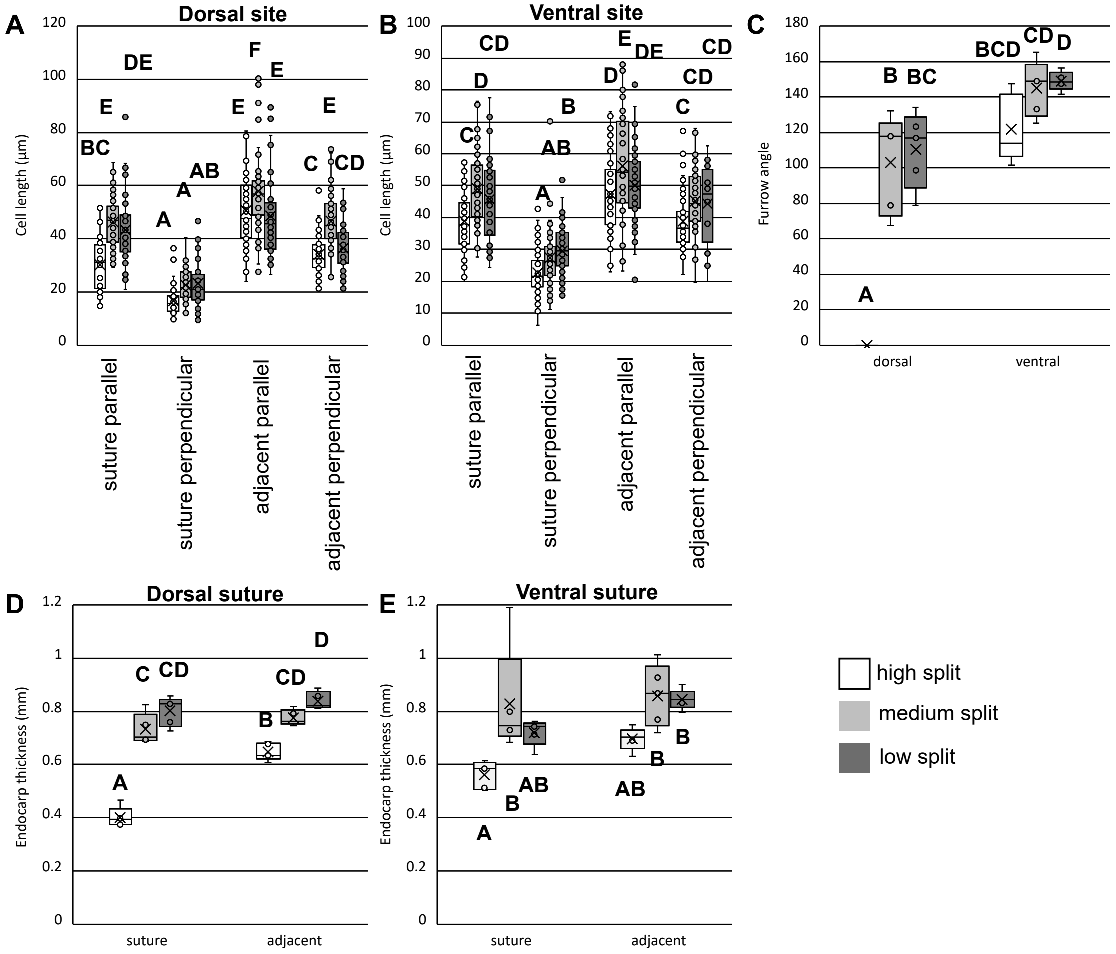

### supplemental figure 5

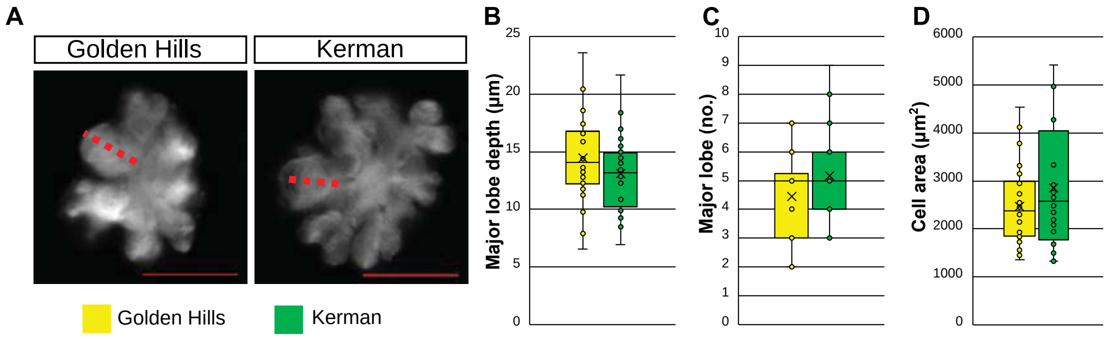
